# 3dcon: tomogram denoising by deconvolution

**DOI:** 10.64898/2026.06.15.732138

**Authors:** Peter Kirchweger, Lev Melnikovsky, Shahar Seifer, Michael Elbaum

## Abstract

Cryo-electron tomography is an expanding technology for the study of macromolecules, viruses, and cells. It is often applied to specimens that are too large or heterogeneous for methods based on 2D image averaging such as single particle analysis, e.g., intracellular membranes or organelles. Current practice records a tilt series of projection images in rotation. Reconstruction is normally an ill-posed mathematical problem. Particularly for the under-determined case of sparse data, discrete tilt angles, and a limited tilt range, characteristic artifacts appear in the reconstructed slices. Much of what appears as noise is in fact structural: the projection of contrast from different planes. Various schemes are employed to regularize the reconstruction, including machine-learning frameworks built on neural networks. To the extent that the noise is structural, it might be suppressed by deconvolution with a suitable kernel. This was demonstrated and has been used regularly in cryo-STEM tomography of thick specimens where the under-sampling problem is particularly acute. Here we present `3dcon` as an open-source extension of the entropy-regularized deconvolution algorithm that had been adopted from fluorescence microscopy. It takes advantage of modern computing hardware for convenient and fast processing. Deconvolution is entirely algorithmic, meaning that successful processing of the data does not depend on the data itself. As such it should be robust in a wide variety of applications.

## 1 Introduction

Tomography is a widely used technique in many fields of science and medicine. It may be defined rather generally as a reconstruction to higher dimension from projections, for example from two-dimensional (2D) images to a 3D volume. For the particular case of projections around a single axis of tilt, the Radon transform describes the transformation from radial to Cartesian coordinates. The inverse transform, from projections to volume, is the essence of tomographic reconstruction. In many cases this operation is ill-posed mathematically due to sparsity of the original data. Therefore the reconstruction requires some form of regularization.

Cryo-electron tomography (cryo-ET, Frank 2006), the case at hand here, is used to visualize unstained biological samples in near-native environments in 3D. For a given dataset size, there is a necessary compromise between resolution and field of view (FOV). In cryo-ET this begins with the microscope configuration. Typically, the wide-field modality of conventional transmission electron microscopy (TEM) is optimal for high resolution, while tomography based on scanning transmission electron microscopy (STEM) is better suited for larger FOV, and in particular for thicker specimens (Elbaum et al. 2021; Varsano and Wolf 2022).

In cryo-ET, acquisition of a tomographic tilt series suffers from a number of characteristic limitations. Foremost, a planar specimen cannot be tilted perpendicular to the illumination, so there exists a large range of inaccessible tilt angles surrounding the surface normal; typically, the rotation is restricted to about −60° to 60°. This creates a large “missing wedge” of information in the upward and downward orientations of the resulting 3D-volume. Furthermore, the projections are normally recorded at discrete tilt angles, due to the requirements of mechanical stability during the recording. The discrete tilt steps lead to smaller missing wedges in between the step directions (Penczek 2010). More subtly, the sensitivity of delicate cryogenic specimens to the ionizing electron irradiation limits the total exposure rather strictly, so that a balance must be struck between the number of tilt views acquired and the dose applied to individual tilt images. The Crowther criterion equates roughly between the number of distinct acquired views and the number of independently determined virtual slices in the reconstruction (Crowther, DeRosier, and Klug 1970). These geometric considerations introduce specific artifacts in the reconstruction, e.g. an elongation in z, anisotropic resolution, and streaking artifacts (most notably on high intensity features). In any given xy plane, much of what appears as random noise is actually a projection of intensity from distant planes. As a structural artifact, it should be possible to suppress this apparent noise based on knowledge of the reconstruction process.

The practical implementation of tomographic reconstruction involves two distinct steps. Alignment of the tilt series compensates for mechanical or optical deviations from the ideal of rigid body rotation around a single axis, or even distortion of the sample itself that may occur during the tilt series (Fernandez and S. Li 2021; Chaillet et al. 2026; Chen 2026). Reconstruction may then be performed by a number of related algorithms. These include most commonly direct back-projection (BP) in real space or Fourier reconstruction and SIRT (or SART), which compares re-projections of the reconstruction to the original dataset and iteratively minimizes the difference. Importantly, the contributions of the tilt views in reciprocal space represent a structure of co-axial tilted planes, so the sampling of spatial frequencies is non-uniform. BP and Fourier reconstruction, for example, result in over-weighting of the low spatial frequencies and a hazy appearance in virtual slices of the reconstructed volume. This is normally compensated by weighting to suppress the low and boost the high frequencies. The choice of weighting function strongly affects the texture of the image, and reconstruction becomes something of an art.

An inherent characteristic of BP is that a bright point in a projection forms a continuous ray in the reconstruction (e.g. Fig. 6 in Carazo et al. 2006). While the projections from multiple views converge to a spot at the point of intersection, the contrast between this intersection and the rays themselves is only on the order of the number of tilts. When viewed at a distant plane in the reconstruction, these rays will appear as a series of spots. Considering that all points in the projections, of any intensity, radiate to rays, the result is a “salt and pepper” noise that permeates the reconstructed volume. The origin of this noise is entirely structural, based on the geometry of the tilt scheme, in contrast to random noise associated with dose limitations or instrumentation. Such a geometrical artifact may be described as a convolution with a kernel, and a means to calculate this kernel was proposed based on sampling theory (Seifer 2023). Note that the reconstructed volume also involves a convolution with the point spread function (PSF) of the physical illumination by the electron beam. Since the kernel is itself a finite array of voxels, it is calculated by accurate integration over the intensity of the scanning beam. These two features are effectively combined into a single kernel by summation of multiple copies of the single beam representing the optical illumination, tilted to the set of angles used in generating the reconstruction. Due to its appearance, we call this kernel the “fanout” PSF.

In a number of earlier works, we have demonstrated that contrast in BP reconstructions is qualitatively improved by 3D deconvolution (Waugh et al. 2020; Croxford et al. 2021; Kirchweger, Mullick, et al. 2023). The kernel represents the idealized BP reconstruction of a bright point under the physical illumination of a particular experimental condition. The most obvious results are noise reduction in 2D virtual sections and a very significant improvement in the axial, i.e., depth, resolution, resulting in greatly enhanced interpretability.

The most successful and stable algorithm in our experience has been the entropy-regularized, iterative deconvolution (ERDC) developed originally for fluorescence microscopy by Muthuvel Arigovindan at UC San Francisco (Arigovindan et al. 2013). It is based on the use of a unique regularization term with intensity-dependent penalization of the second order derivatives. This approach manages to suppress noise effectively while at the same time preserving essential details, and the processing is robust to noise even after many iterations. Unfortunately, the code is not open source nor actively maintained, and sustainability has become a major concern. Execution might also be accelerated and accessible volume expanded by taking advantage of modern advances in computer architecture such as GPU. Moreover, our established workflow combines several different software environments that reflect more heavily the history of the development than proper requirements for implementation. Therefore we undertook a rewriting of the code from scratch, inspired by the original publication but geared more specifically to the tomography application. We expect that the resulting software presented here should be accessible and conveniently applicable by the community to 3D reconstruction originating from both STEM and TEM modalities.

## 2 Application

In order to demonstrate the application of `3dcon` we used a previously published dual-axis cryo-STET (data to be found on [Zenodo](https://doi.org/10.5281/zenodo.14959968) and the EM-databank EMD-51764, Kirchweger, Wolf, et al. 2025) dataset of adherent mouse cells from our lab (Fig 2) and a well-known TEM-based cryo-ET (EMPIAR-10164, Schur et al. 2016) dataset of HIV-1 virus-like particles (VLP) (Fig 3). Processing begins with a simple (unweighted) BP reconstruction, i.e., after the series alignment as might be performed in IMOD or AreTomo. A fanout PSF is then generated with a beam profile appropriate to the angles used in BP. (Note that these may differ from the precise goniometer angles if optimized in alignment, viz., the .tlt vs. the .rawtlt file in IMOD.) Deconvolution is then executed as an iterative procedure. In each case, we aimed for `3dcon` settings that produce high-quality images at a small number of iterations (e.g. around 20) while remaining stable over a much larger number (200 ∼ 500).

**Figure 1.**
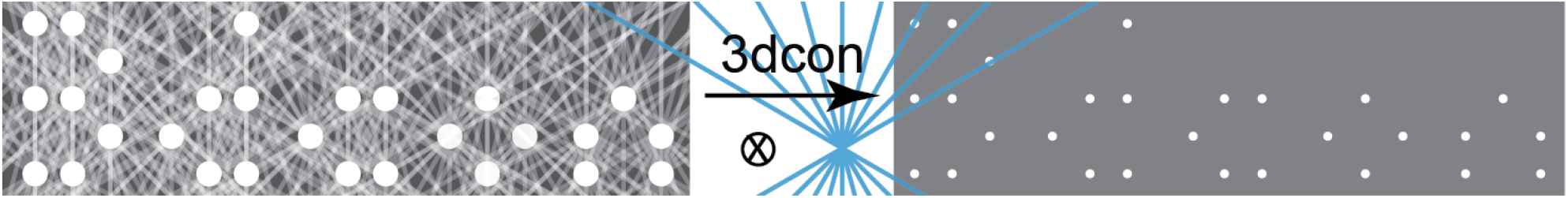
Schematic representation of the `3dcon` improvements. A slab-like sample permits recording of projection images over a limited angular range, resulting in a large missing wedge of information at top and bottom. In addition, recording of projections at discrete angular intervals results in as many small missing wedges in between. Deconvolving the 3D volume in XZ-direction with a fanout kernel (blue) mimicking the tilt directions maps the streaks in the reconstruction back to their origin, thereby improving the reconstruction quality.

**Figure 2.**
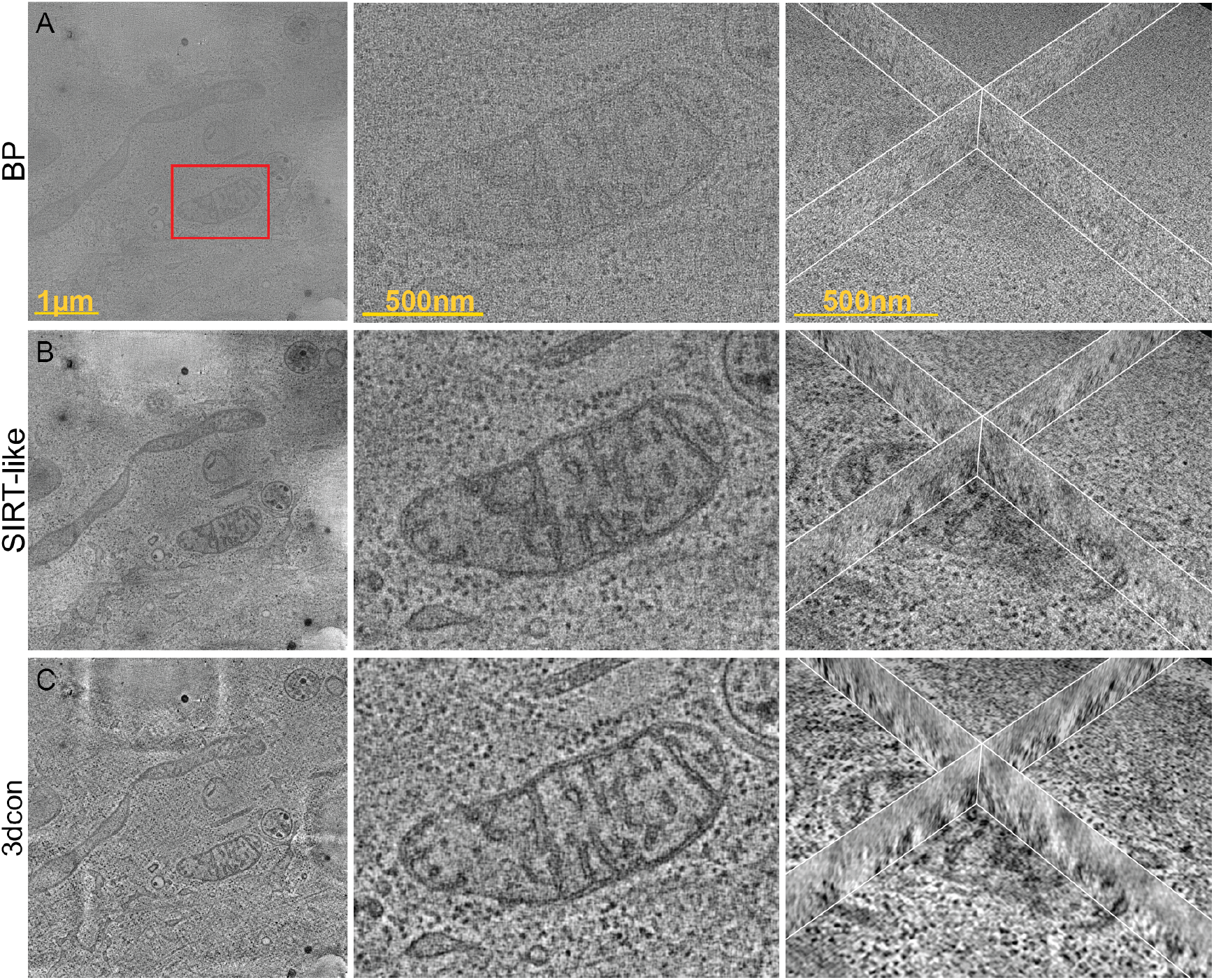
Comparison of (A) BP, (B) SIRT-like, (C) `3dcon` of a dual-axis cryo-STET data-set (EMD-51764, Kirchweger, Wolf, et al. 2025). 1st column show individual slices of the whole tomogram (scale: 1 µm). 2nd column zoom into one the mitochondria, as highlighted by the red box (scale: 500 nm). 3rd column shows orthoslices (scale: 500 nm).

**Figure 3.**
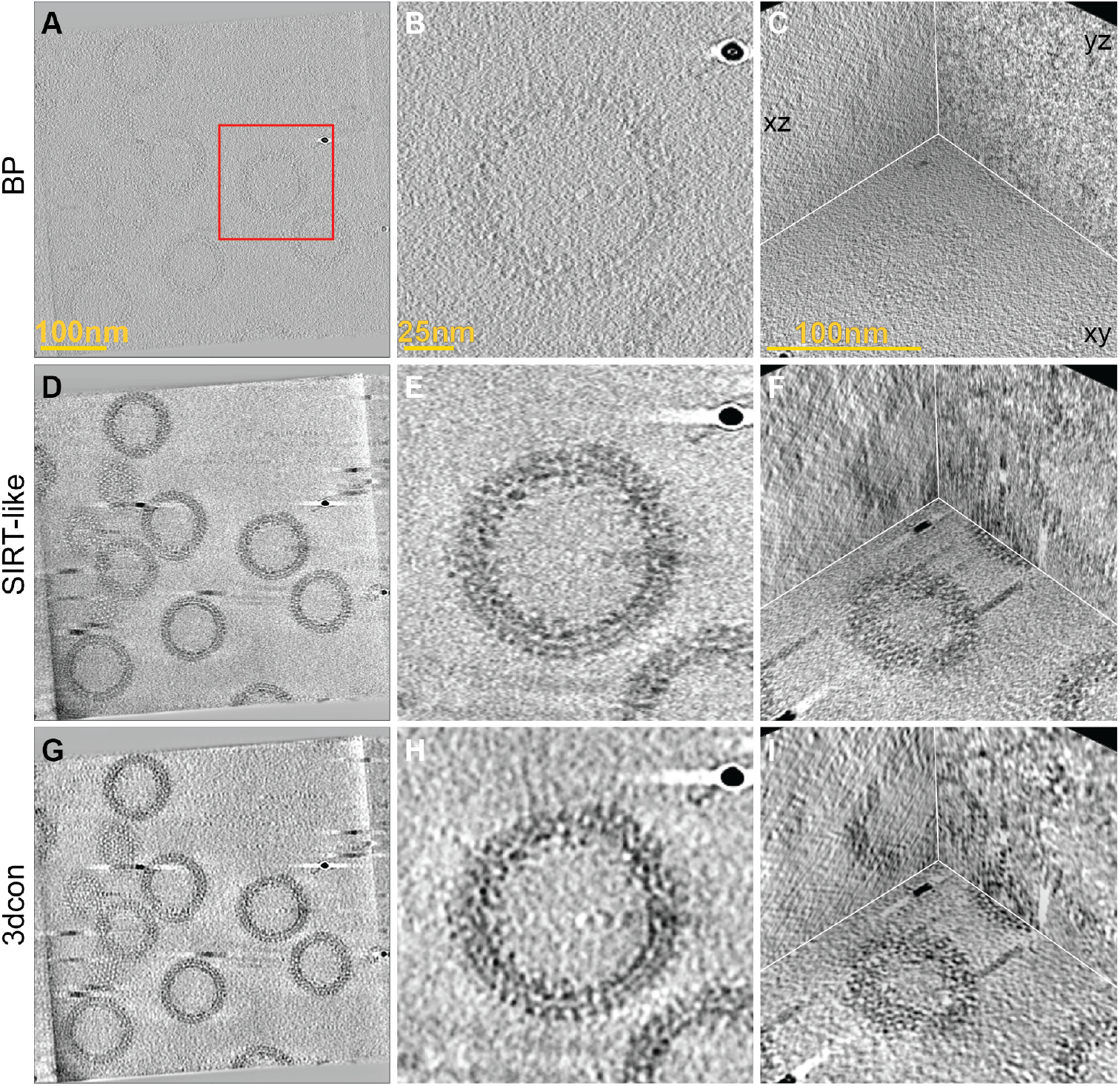
A cryo-TEM tomogram (EMPIAR-10164, Schur et al. 2016) processed with `3dcon`. (A,B,C) show the BP, (D,E,F) the SIRT-like filtered, and (G,H,I) the `3dcon` of a slice in the tomogram (first column, scale: 100 nm), a VLP (middle column, scale: 25 nm) highlighted by the red box and orthoslices through a VLP (scale: 100 nm).

**Figure 4.**
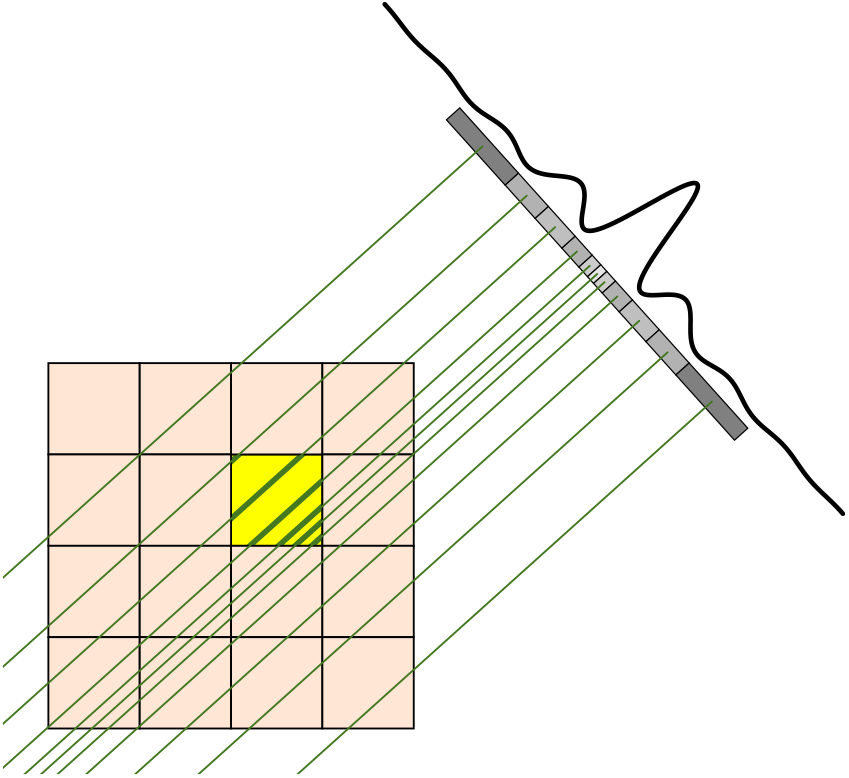
Schematic representation of the PSF rendering algorithm. The contributions of each ray for all beam directions are summed to produce the final PSF voxel array.

Dual-axis tomography reduces the large missing wedge to a missing cone, resulting in a similar appearance for XZ and YZ orthosections of the reconstruction. As such, it affords a good opportunity to focus on the denoising effect of deconvolution on the small missing wedge artifacts. The data were acquired using a workflow that we described recently for microscopes equipped with a dual-axis stage (Kirchweger, Wolf, et al. 2025); the SerialEM script automatically returns to the same list of stage locations (Navigator points) after rotation of the grid by 90 deg. The reconstruction presented in Fig 2 displays an unweighted BP, a SIRT-like filter, and processing by `3dcon` with 20 iterations. In the XY view (middle column), protein densities decorating the cristae are resolved here in a sample 550 nm thick (Fig 2C). The improvement afforded by `3dcon` is most apparent in the orthoslices (right column), where image sharpness is significantly improved. (For completeness, we show the final 200 iterations, and a `3dcon`-run with an additional smoothing factor in Fig S1). For comparison, conventional reconstructions from the first-acquired “a”-axis tomogram appear in Fig S2. Again, `3dcon` improves upon the SIRT-like reconstruction, and the display of a result after 200 iterations is intended to emphasize the stability of the algorithm.

To demonstrate that `3dcon` can also be applied to TEM-based cryo-ET data, we followed the pipeline in nextPYP (H.-F. Liu et al. 2023) for processing of virus-like particles (VLP) from HIV-1 (EMPIAR-10164, Schur et al. 2016). We generated a contrast transfer function (CTF)-corrected reconstruction by unweighted BP, generated a fanout PSF using the reported tilt angles, and then ran a deconvolution using `3dcon`. Fig 3 shows the improvement of `3dcon` starting from BP by comparing XY planes and orthoplanes of the tomogram. In the BP tomogram (Fig 3A-C), the outline of the VLP is clearly visible, but details are hard to see. The SIRT-like filter improves the contrast very substantially (Fig 3D-F). However, only `3dcon` (Fig 3G-I) recovers most clearly the CA-SP1 hexamer on the VLP. As an estimate of the improvement due to `3dcon`, we calculated the Fourier Shell Correlation (FSC) from the BP, SIRTlike and `3dcon` tomograms, with curves shown in Fig S3A. The curves give an impression that the resolution of the SIRT-like filtered tomogram (orange) is higher at the 0.5 threshold than that of the `3dcon` (green). When inspecting a low-pass filtered zoom-in onto the top of a VLP, however, the contrast in the SIRT-like filtered tomogram (Fig S3B) is less interpretable than that of the `3dcon` (Fig S3C). For completeness, and to show that `3dcon` remains stable over a large number of iterations, we show the same `3dcon` settings with 500 iterations (Fig S4A), and in addition with a smaller smoothing factor (Fig S4B+C).

Finally, we compare `3dcon` to existing tools for denoising (Fig S5). Specifically, for the dual-axis cell tomogram we compare `3dcon` with ERDC (Fig S1D). We find that `3dcon` performs at least as well as ERDC, as expected. Additionally, we compare `3dcon` with CryoCARE (Buchholz, Jordan, et al. 2019; Buchholz, Krull, et al. 2019) and with Isonet2 (Y.-T. Liu et al. 2025). To this end we first split the a-axis of the cell tomogram to even vs odd tilts. CryoCARE produces a visually-pleasing result, albeit smoother than that of `3dcon`. Isonet2 produces very strong contrast at membranes but areas within the mitochondrial matrix appear to be smeared. Possibly this could be improved by fine-tuning of internal parameters such as the box size for network training. For the VLPs, both CryoCARE and Isonet2 appear to be more effective than `3dcon` in quieting the background. However, certain artifacts appear in the particles themselves, most notably features of inverted (white) contrast within the protein shell. `3dcon` should be robust against such features that may be introduced by methods based on deep learning. The underlying architecture of Isonet and its very impressive performance in improving axial resolution for the VLPs suggest that the U-Net filter generated from numerous sub-tomograms might partly represent a 3D PSF. Axial resolution in the case of the thicker mitochondrial tomogram is much less impressive, however. The training to recognize the missing wedge and develop a suitable filter come from the dataset itself. This might explain the variable performance in these two examples. Execution time of `3dcon` is significantly faster than the other methods, as detailed in Tab 1.

**Table 1.** Execution times.

| Program | Hardware | Process | Processing time |
| --- | --- | --- | --- |
| Tomogram size: 2048x2048x238 pixels (Fig 2) |  |  |  |
| ERDC | 48 CPUs<br>256 GiB RAM | fanout generation | 75 min |
|  |  | core2decon | 89 min |
| 3dcon<br>(20 iter) | 2*(NVIDIA L40S)GPUs<br>24 CPU cores<br>70 GiB RAM | fanout generation | 2 min |
|  |  | 3dcon | 8 min |
| CryoCARE | 2*(NVIDIA L40S)GPUs<br>32 CPU cores<br>128 GiB RAM |  | 43 min |
| IsoNet2 | 4*(NVIDIA L40S)GPUs<br>40 CPU cores<br>128 GiB RAM |  | 230 min |
| Tomogram size: 648x648x341 pixels (Fig 3) |  |  |  |
| ERDC | 48 CPUs<br>256 GiB RAM | fanout generation | 5 min |
|  |  | core2decon | 8 min |
| 3dcon<br>(20 iter) | 2*(NVIDIA L40S)GPUs<br>8 CPU cores<br>24 GiB RAM | fanout generation | 0.2 min |
|  |  | 3dcon | 1 min |
|  | Apple MacBook Air<br>M2: 8 CPU cores<br>24 GiB RAM | fanout generation | 0.2 min |
|  |  | 3dcon | 17 min |
| CryoCARE | 2*(NVIDIA L40S)GPUs<br>32 CPU cores<br>64 GiB RAM |  | 38 min |
| IsoNet2 | 2*(NVIDIA L40S)GPUs<br>32 CPU cores<br>64 GiB RAM | | 190 min ($\div 5$) |

In order to compare the execution times on similar hardware, we limited IsoNet2 and CryoCARE to 2 GPUs. We note that execution time of IsoNet2 was the same when processing 1 or 5 tomograms in parallel.

## 3 Implementation

Deconvolution aims to invert the effect of the convolution between an object *g*(**r**) and a kernel *h*(**r**) that occurs during data acquisition, resulting in the measured data *f*(**r**):

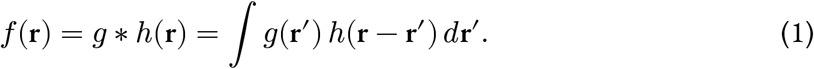

It is essential that the kernel depends only on the relative positions **r** − **r**′. Not all processes satisfy this condition. Important examples that can be approximated as convolutions include fixed out-of-focus purely geometric blurring, constant velocity motion blur, parallel beam tomography followed by simple BP, and diffraction-limited imaging in the weak phase limit. Since a composition of multiple convolutions is also a convolution, a single deconvolution operation can in principle invert them all simultaneously.

Artifacts in data acquisition for STEM tomography arise from both the diffraction-limited probe and the BP reconstruction. Due to the finite convergence angle *α* of the focused illumination, the former can be approximated as a simple convolution for thin specimens:

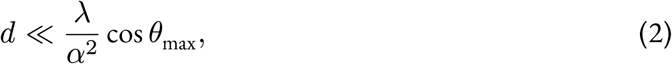

where *d* is the specimen thickness, *θ*_max_ is the maximum tilt angle, and *λ* is the electron wavelength. Similarly, for TEM tomography, where the significant defocus Δ*f* required to generate adequate contrast is also responsible for the image blurring, the specimen thickness is also limited:

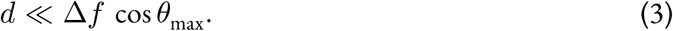

Deconvolution is traditionally formulated as a minimization problem

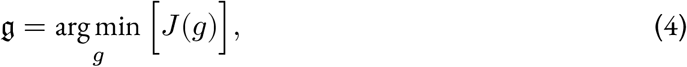

where *J*(*g*) is a loss function that quantifies the quality of the result *g*(**r**) given the input data and prior knowledge. The loss function suggested in (Arigovindan et al. 2013) consists of three terms

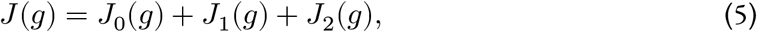

where the regularization *J*_1_ and positivity *J*_2_ terms impose prior knowledge about the object, while *J*_0_ enforces agreement with the input data. The latter is estimated as the mean-squares difference

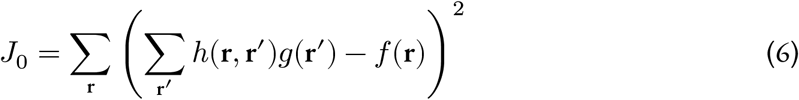

between the input data *f*(**r**) and the result convolved with the kernel *h*(**r**). This kernel, also known as the point spread function (PSF), in the case of STEM tomography reconstruction is the fan-shaped response to a point source after BP. For convenience, we built two separate tools: (i) `fancy` — to generate the PSF, and (ii) `3dcon` — to actually perform the deconvolution. Alternatively, the deconvolution may be performed using an externally generated PSF.

### 3.1 PSF generation with fancy

The diffraction-limited probe in STEM is approximated by a parallel beam with an Airy disk intensity profile:

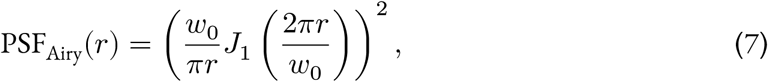

where *J*_1_ is the Bessel function of the first kind, *r* is the radial distance from the center of the probe, and *w*_0_ is given by

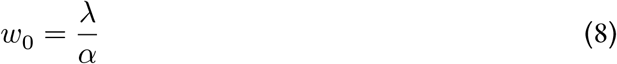

with *λ* being the electron wavelength and *α* the convergence semi-angle. The complete PSF is a superposition of such probes for all beam directions, i.e., for all tilt angles. Of course, STEM uses a focused illumination; the parallel beam condition (2) expresses the presumption that the specimen is thinner than the depth of focus. Alternatively, if the depth-dependent magnification is taken into account, a 4D STEM image may be computationally refocused by techniques based on parallax (e.g., tcBF, Yu et al. 2025), synchronization (shadow montage, Seifer, Houben, and Elbaum 2026), or ptychography using a deliberately defocused probe (Varnavides et al. 2026).

For an accurate deconvolution, the PSF must be represented as a 3D voxel array at least twice the size of the input volume in each direction. Each voxel is assigned the integrated PSF intensity, which presents major implementation difficulties because of the array size. This is partially alleviated by the fact that the beams are sparse, and contribution of the parts beyond a few units of *w*_0_ can be neglected. Nevertheless, an efficient integration algorithm is needed to compute the PSF in reasonable time. This is achieved by the following protocol:

1. numerically pre-computing integrals of (7) on an adaptive fine 1D partition;
2. generating a 2D polar grid using the partition from the previous step;
3. assigning integrated profile values to each 2D grid cell;
4. approximating full beam as a superposition of straight rays emitted from the centers of each 2D grid cell;
5. tracing each ray in space and summing their contributions to the 3D PSF voxel array;
6. repeating the previous two steps for all beam directions.

This approach, as implemented in `fancy`, allows arbitrary beam orientations, such as those used in dual-axis tomography or single-particle analysis, and provides the flexibility to accommodate other beam profiles than the simple Airy disk.

### 3.2 Deconvolution with 3dcon

We largely follow the notation of (Arigovindan et al. 2013), where ERDC software is introduced. We emphasize here only the important implementation decisions that are not explicitly detailed in that work, or where our approach differs from the original paper.

ERDC stands out by its use of the entropy-based regularization term *J*_1_, which contains second spatial derivatives of the reconstruction *∂*^2^*g*/*∂x*^*i*^*∂x*^*j*^ to penalize high spatial frequency components. One can build two quadratic scalars from these derivatives: the sum of squares of all second derivatives, and the square of the Laplacian. We combine both scalars in our implementation:

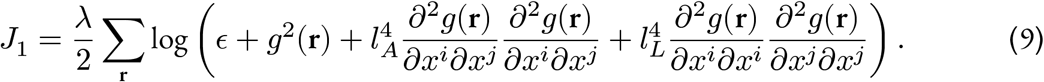

Characteristic lengths for the two types of smoothing *l*_*A*_ and *l*_*L*_ can be set independently and specified either in voxel dimensions or in Angstroms. Parameter *λ* controls the overall strength of the regularization, while *ϵ* is adjusted to tune the contributions of intensity and smoothing terms. We use symmetric stencil finite differences to compute all derivatives in real space. For completeness, we also explicitly write down the positivity constraint term

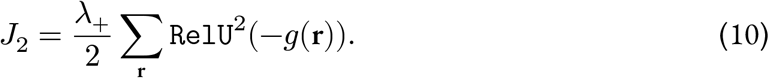

All code is implemented in Python, utilizing flexible CPU/GPU vectorized operation via `PyTorch` (Ansel et al. 2024) wherever the computation speed is critical. Other parts rely on the widely used `numpy` library (Harris et al. 2020).

Depending on available hardware, the code allows for parallel execution and features fast execution even on large volumes. This is exemplified in Tab 1, where we compare ERDC (on a cluster node with 48 CPUs) with `3dcon` (on a node with 2 GPUs, 8 CPUs and 24 GiB RAM, and a MacBook Air with 8 M2 CPUs and 24 GiB RAM).

Special care is taken to reduce memory consumption by optionally splitting the volume into chunks. Although the PSF itself is not separable, the Fourier transform is separable, allowing us to perform convolution sequentially. This approach also enables utilization of GPUs with limited memory or effective use of CPU cache. There is no hard upper limit to the tomogram size as had been the case for ERDC. However, `3dcon` runs of tomograms that do not fit into local memory significantly prolong running times.

The loss function minimization is performed via robust Conjugate Gradient method. We used a “monkey-patched” `scipy.optimize.fmin_cg` function from the `scipy` library (Virtanen et al. 2020) to reduce memory consumption by avoiding unnecessary copies of large arrays. The variations of all three terms of the loss function with respect to the reconstruction *g*(**r**) are computed analytically

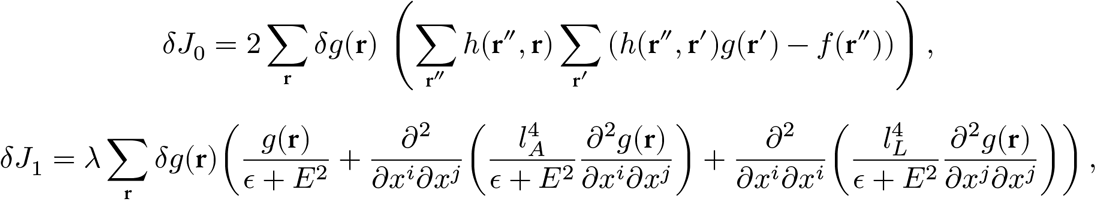

where *E*^2^ is the sum of the last three terms under the logarithm in (9), and

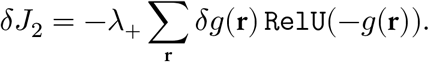

The package is open source for wide usage and flexible for extension to other regularization terms or use in other projects as a library. It is available at https://github.com/elbaum-lab/3dcon.

### 3.3 Tomogram alignment and reconstruction

The dual-axis cryo-STET dataset was aligned in IMOD (Kremer, Mastronarde, and McIntosh 1996; Mastronarde 1997; Mastronarde and Held 2017) as described before (EMD-51764, Kirchweger, Wolf, et al. 2025).

The HIV-capsid dataset was described before (EMPIAR-10164, Schur et al. 2016). The dataset was imported to nextPYP (H.-F. Liu et al. 2023) and aligned using fiducials. Tomograms from even and odd tilt series were produced as well.

As a starting model for `3dcon` we used unweighted back projection tomograms. This were produced in IMOD by setting the flag RADIAL to 0.5 0.5.

### 3.4 Fanout generation

The fanouts were generated using the bundled `fancy` program.

### 3.5 Deconvolution runs

To execute `3dcon` runs, we set the two types of smoothing *l*_*A*_ and *l*_*L*_ to 1 pixel and adapted *λ* and *ϵ*. `3dcon` settings in the single-axis STEM example shown here included *λ* set to 0.001 and *ϵ* to 0.0001 in Fig S1C-D (20 and 200 iterations, respectively). `3dcon` settings in the dual-axis STEM example shown here included *λ* set to 0.001 and *ϵ* to 0.0001 (Fig 2C and Fig S2B, 22 and 200 iterations, respectively), and *λ* set to 0.001 and *ϵ* to 0.0001 (Fig S2C,D, 21 and 200 iterations, respectively). In the TEM example, settings included *λ* set to 0.001 and *ϵ* to 0.0001 (Fig 3G-I and Fig S4A, 23 and 500 iterations, respectively), and *λ* set to 0.01 and *ϵ* to 0.0001 (Fig S4B,C, 21 and 500 iterations, respectively).

### 3.6 Visualization

All tomograms were visualized in ChimeraX (Pettersen et al. 2021), except for FigS4 which were saved from 3dmod (Kremer, Mastronarde, and McIntosh 1996).

### 3.7 FSC calculation

To calculate the FSC, a python code was written, which estimates the resolution in a 2D projection or 3D volume on a checkerboard-based pattern. This FSC-calculation script is found on GitHub together with a Jupyter notebook showing how to use it,https://github.com/elbaum-lab/fsc.

## 4 Discussion

3D deconvolution is an effective approach to ameliorate the reconstruction artifacts of BP from a discrete set of view angles (Waugh et al. 2020; Kirchweger, Mullick, et al. 2023; Seifer 2023). While these artifacts are recognized as distortions that appear in orthoslices along the untilted view direction (i.e., xy or xz), they are also a primary cause of noise in virtual sections throughout the depth (i.e., cuts in xy). It should be emphasized that the relevant noise is structural, inherent in the interaction of the illumination with the sample, rather than statistical due to low electron count or poor detector efficiency. As such, the standard Poissonian or Gaussian noise models are suitable by coincidence at best. Deconvolution should in principle be an integral step in reconstruction by BP. To date, 3D deconvolution has been applied mainly to cryo-STEM tomography, where the artifacts are exaggerated by sample thickness (Elbaum 2018; Waugh et al. 2020; Elbaum et al. 2021; Kirchweger, Mullick, et al. 2023; Kirchweger, Seifer, et al. 2025; Kirchweger, Wolf, et al. 2025) and the ER (entropy regularized) algorithm (Arigovindan et al. 2013; Seifer 2023) was found to work most effectively. A similar approach was used for dual-axis tomography, resulting in near-isotropic resolution across the orthoslices (Kirchweger, Mullick, et al. 2023). Use of ER decon has also been demonstrated for wide-field TEM tomography (Croxford et al. 2021). Recently, a similar approach was employed for molecular data processed by single particle analysis. There it was shown that deconvolution reduces anisotropy in the reconstruction that originates from preferred orientations (J. Li et al. 2025). We should emphasize that ERDC is a distinct operation from that of contrast transfer function (CTF) correction or low-frequency contrast enhancement as offered in Warp (Tegunov and Cramer 2019), both of which rely on a similar mathematical operation but implement differently and use a different kernel.

The goal of this work has been to adapt the original ERDC software for electron tomography and to modernize it in light of subsequent developments in computer hardware. `3dcon` is written in Python and distributed as open source software under the GPLv3 license for sustainability. While maintaining the original features, several specializations and extensions have been added. In particular, the package includes a second regularization option based on the square of the Laplacian and a convenient PSF generator for tilt tomography. The latter appears as a fanout rather than a radially-symmetric cone as in fluorescence microscopy. The code can take advantage of thin, nearly planar fanouts to reduce memory requirements and accelerate execution but it is also ready to work with space-filling kernels (e.g., in dual-axis configurations.)

The algorithm was specifically optimized for memory efficiency, allowing efficient CPU cache usage and GPU utilization when available. By taking advantage of modern Python libraries, the code is flexible with respect to hardware, running on multiple CPUs and up to 2 GPUs. Although a server is recommended, `3dcon` runs even on a laptop computer. A very practical aspect of the improved running speed is the ability to rapidly explore the parameter space of the deconvolution. This process frequently requires multiple attempts and can significantly improve the final result.

As one of the benefits of `3dcon` is denoising, we compared the results to cryo-CARE (Buchholz, Jordan, et al. 2019; Buchholz, Krull, et al. 2019) and IsoNet2 (Y.-T. Liu et al. 2025), popular tools in the TEM-based cryo-ET workflow. Cryo-CARE uses a neural net to compare half-datasets in order to isolate and suppress noise on the assumption of statistical independence between the halves. IsoNet2 also uses half-datasets and employs a subvolume averaging approach to learn about the noise in the tomogram. In STEM tomography we do not acquire multiple movie frames per tilt image, so two tomograms were reconstructed from the even and odd tilts. The resulting noise reduction was impressive, especially considering that the underlying noise model is not fully justified. However, we found that the `3dcon` processing better preserves the finer details, and the execution speed is relatively fast. Moreover, it takes a purely algorithmic approach that does not depend on training generated from the recorded data per se. Therefore it is likely to work even when neural networks may not. Provided that the PSF is an accurate description of the acquisition conditions, `3dcon` should provide a precise regularization for a given set of adjustable weights and iterations. We expect that it will become a useful instrument in the tool-chest for electron tomography.

## 5 Acknowledgments

M.E. acknowledges insightful discussions with John Sedat. Computational work was carried out on the Faculty of Chemistry’s high-performance computing facility CHEMFARM at the Weizmann Institute of Science, which is supported in part by the Ben May Center for Chemical Theory and Computation. This work was supported in part with funding from the European Union (ERC-Adv CryoSTEM, 101055413; Views and opinions expressed are however those of the authors only and do not necessarily reflect those of the European Union or the European Research Council. Neither the European Union nor the granting authority can be held responsible for them.) ME is incumbent of the Sam and Ayala Zacks Professorial Chair in Chemistry. This research is made possible in part by the historical generosity of the Harold Perlman family.

