## Supplemental Data for "3dcon: tomogram denoising by deconvolution"

### Supplementary Material
### Advancement of 3D deconvolution in cryo-ET

Peter Kirchweger [1,†], Lev Melnikovsky [1,†], Shahar Seifer [1], and Michael Elbaum [1,*]

[1] *Department of Chemical and Biological Physics, Weizmann Institute of Science, 7610001 Rehovot, Israel*
[†] *These authors contributed equally*

June 14, 2026

### 1 Supplementary Figures

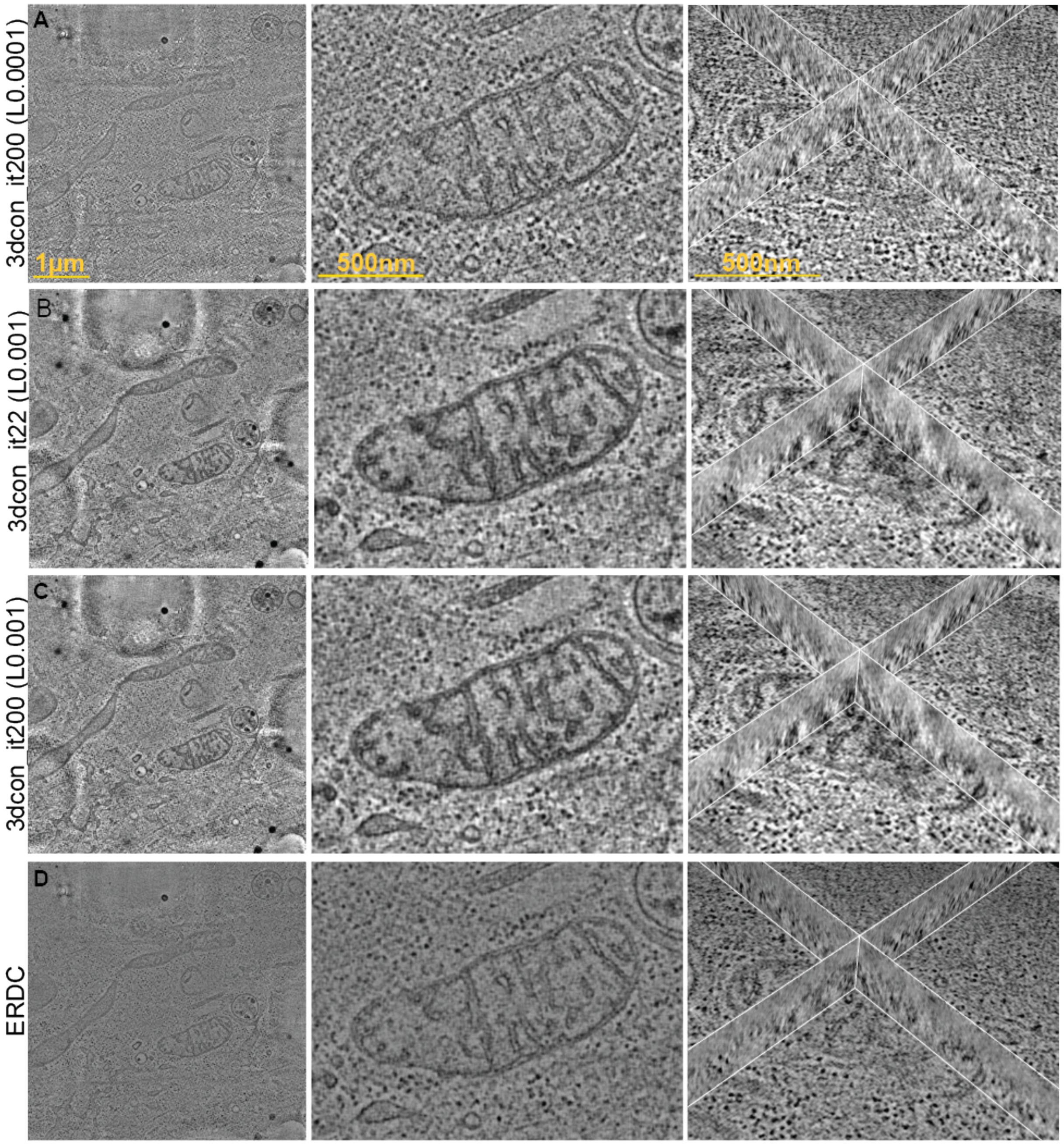


**Figure S1:** Comparison of different iterations and smoothing factors of a `3dcon` (first three rows) and ERDC (bottom row) of the dual-axis cryo-STET dataset (EMD-51764, Kirchweger et al. 2025). 1st column show individual slices of the whole tomogram (scale: 1 µm). 2nd column zoom into one the mitochondria, as highlighted by the red box (scale: 500 nm). 3rd column shows orthoslices (scale: 500 nm).

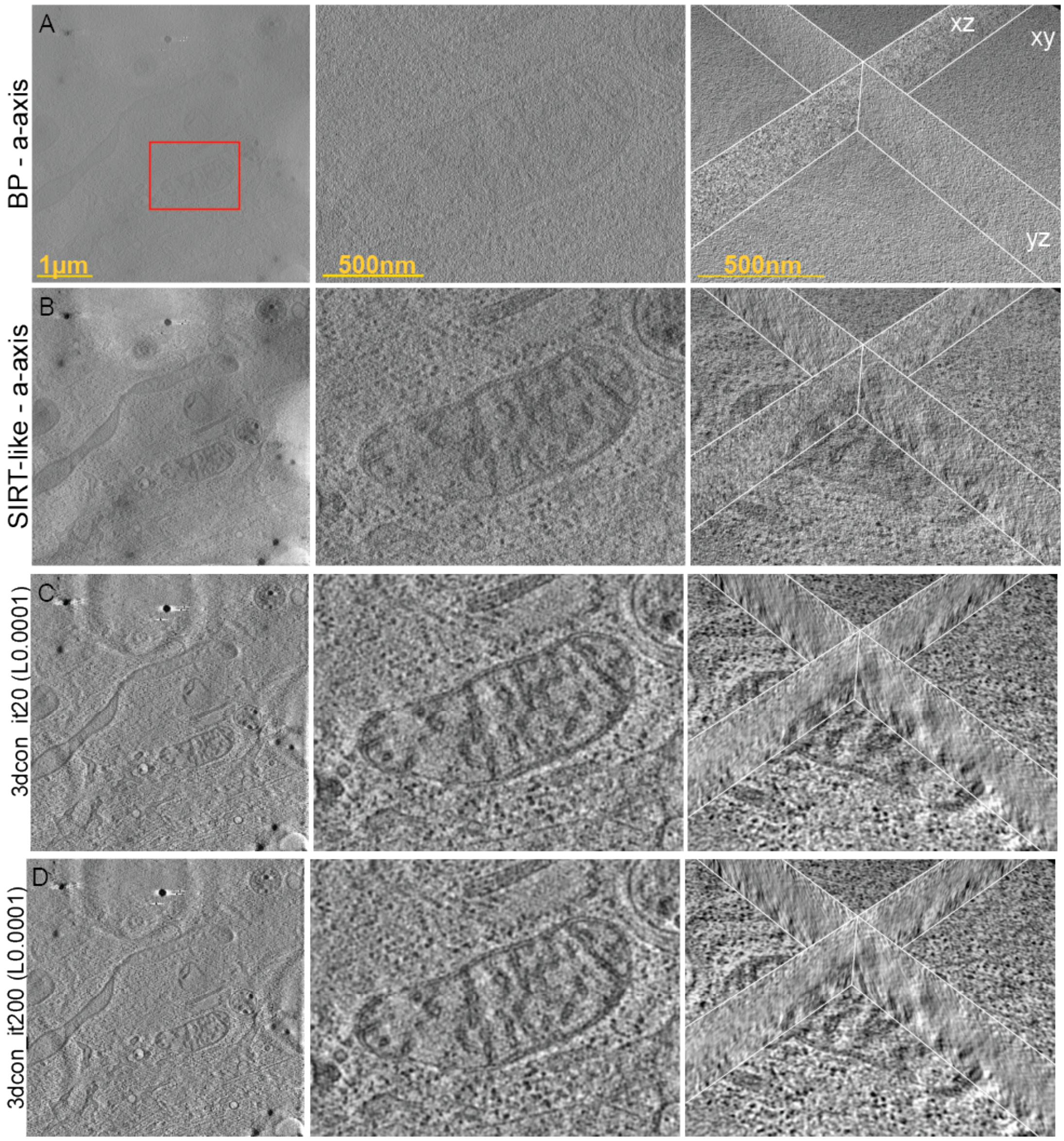


**Figure S2:** Comparison of (A) BP, (B) SIRT-like, (C) `3dcon` (20 iterations) , and (D) `3dcon` (200 iterations) of the a-axis cryo-STET dataset from (EMD-51764, Kirchweger et al. 2025). 1st column show individual slices of the whole tomogram (scale: 1 μm). 2nd column zoom into one the mitochondria, as highlighted by the red box (scale: 500 nm). 3rd column shows orthoslices (scale: 500 nm).

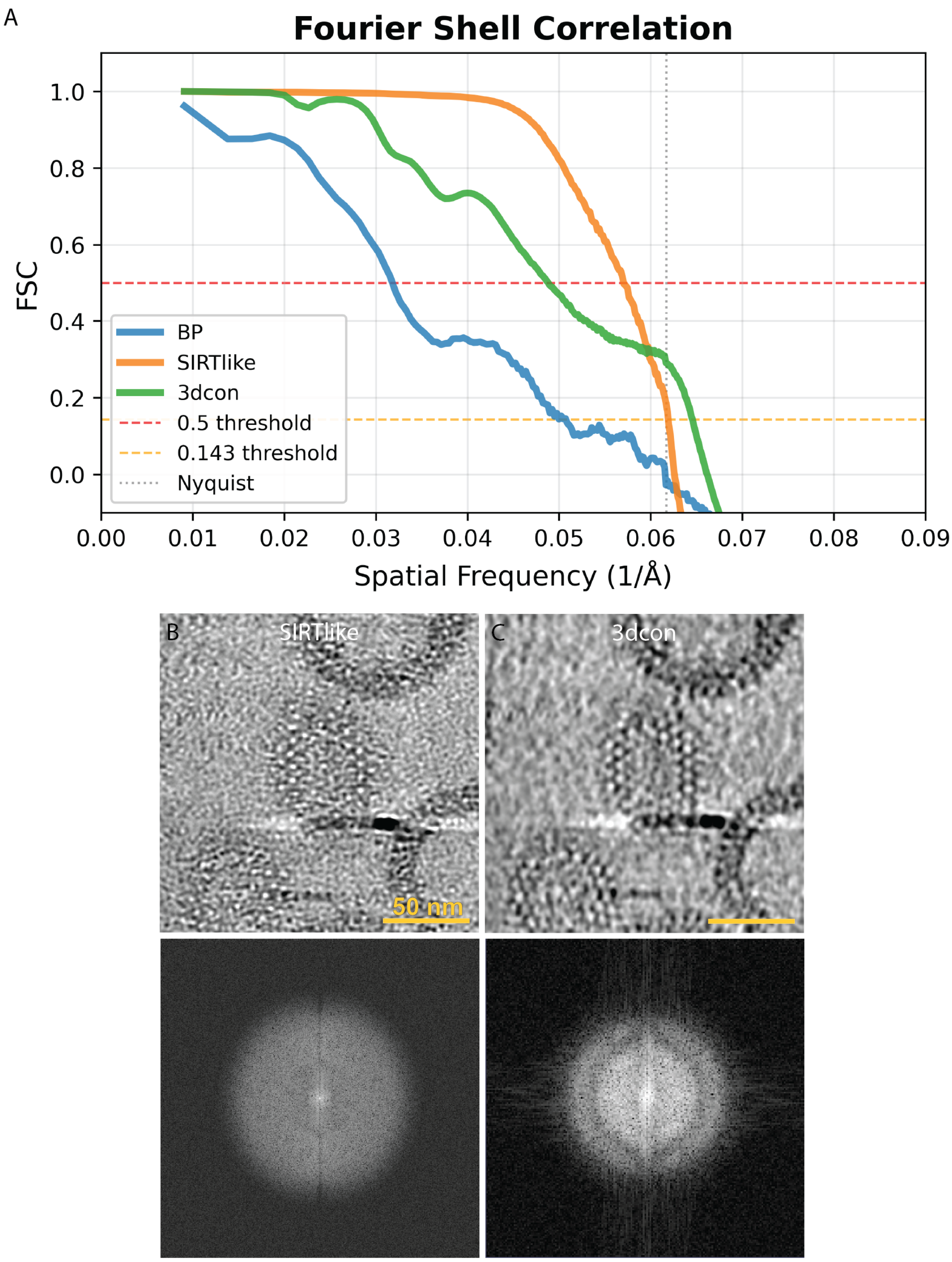


**Figure S3:** Resolution estimates. (A) FSCs from the BP (blue), SIRTlike (orange) and `3dcon` (green) of the cryo-TEM dataset. (B-C) Single slice (top row) and snapshot of the FFT (bottom row) of a low-pass filtered tomogram of (A) the SIRTlike and (B) `3dcon`. The radius for low-pass filtering was set to 0.2. Note that protein density must appear darker than the water background. Scale bar is 50 nm.

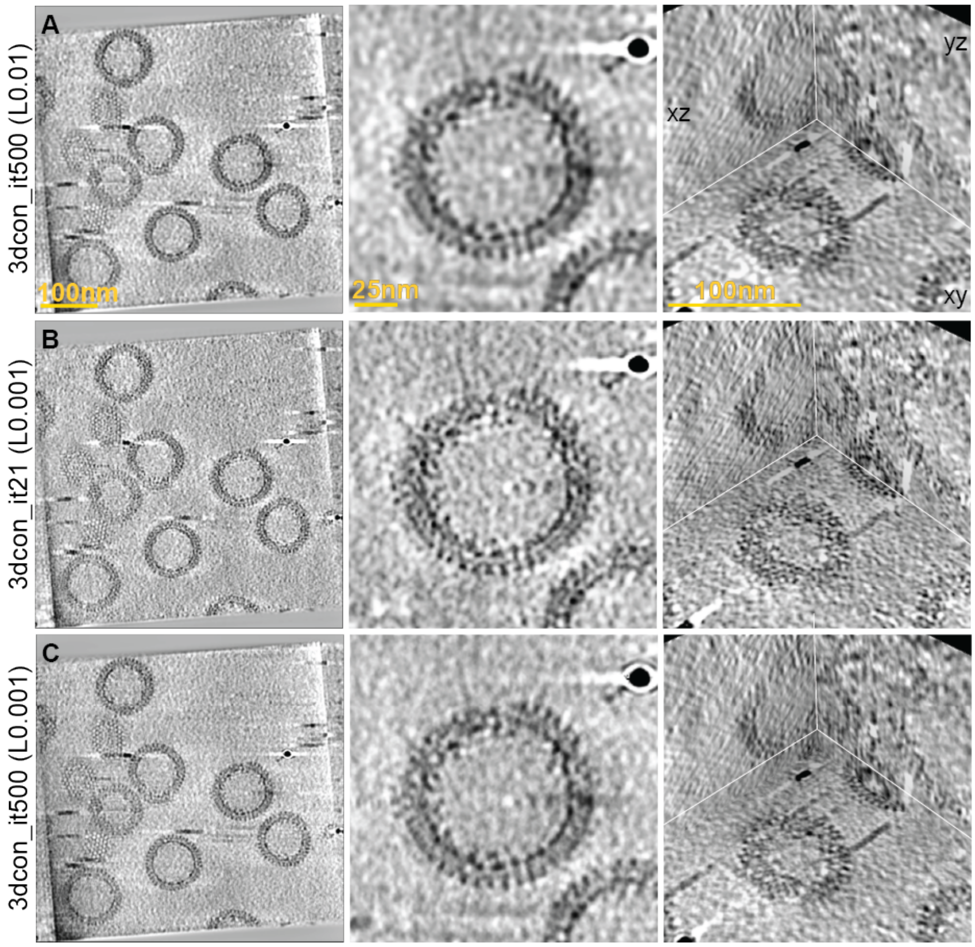


**Figure S4:** Comparison of different iterations and smoothing factors of a `3dcon` (A-C) of the cryo-TEM dataset (EMPIAR-10164, Schur et al. 2016). 1st column show individual slices of the whole tomogram (scale: 100 nm). 2nd column zoom into one the VLP (scale: 25 nm). 3rd column shows orthoslices (scale: 100 nm).

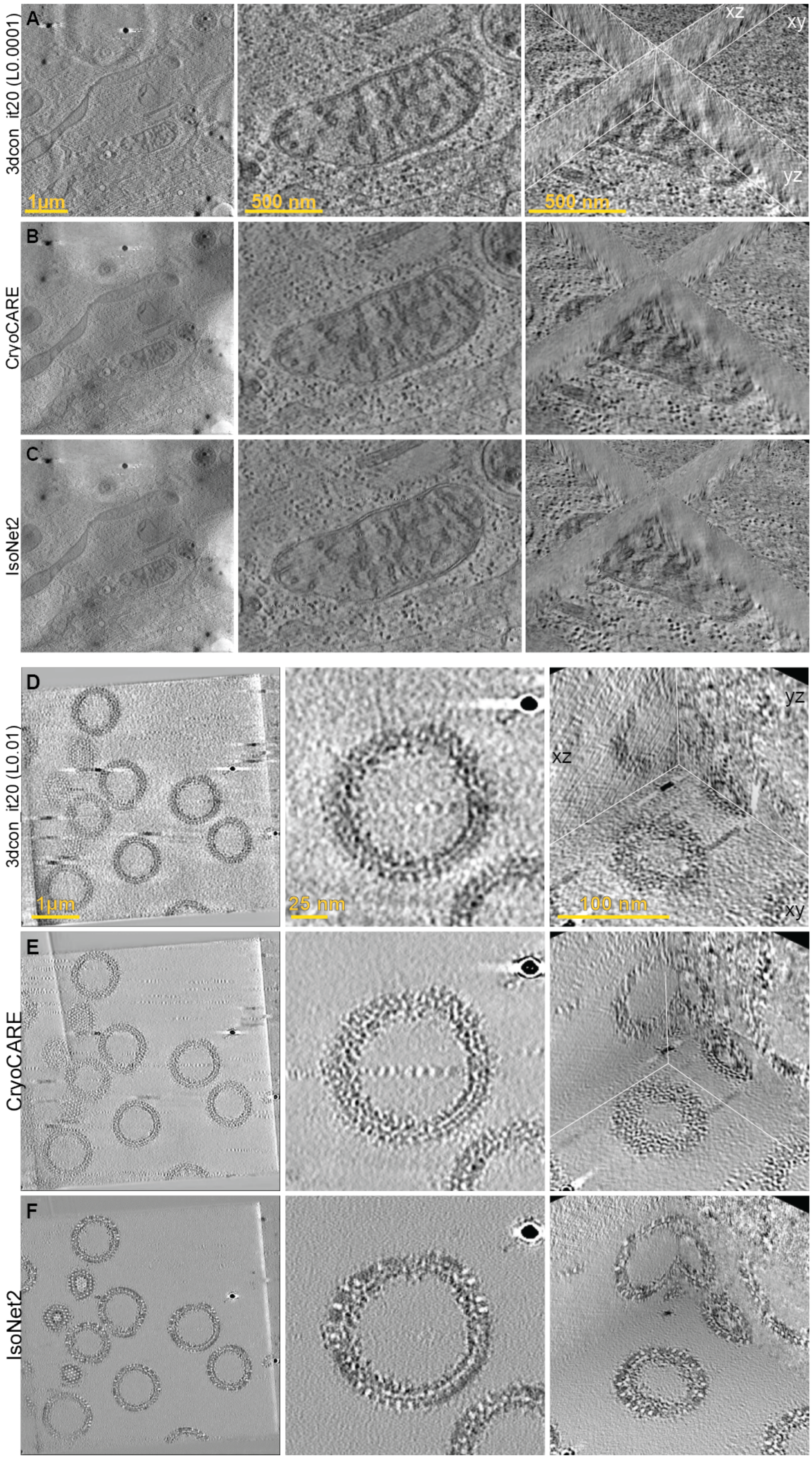


**Figure S5:** Comparison of `3dcon` (A,C) to CryoCARE (B-D) and IsoNET2 (E) of the a-axis of the cell tomogram (A-B) and the VLPs (C-E). 1st column show individual slices of the whole tomogram (scale: 1 µm (A-B) and 100 nm (C-E)). 2nd column zoom into one area (scale: (A-B) 500 nm and (C-E) 25 nm). 3rd column shows orthoslices (scale: (A-B) 500 nm and (C-E) 100 nm). (A,C) are the same images as in Fig 2C and Fig 3G-I, respectively.
